# WormSwin: Instance segmentation of C. elegans using vision transformer

**DOI:** 10.1101/2023.04.10.536324

**Authors:** Maurice Deserno, Katarzyna Bozek

## Abstract

The possibility to extract motion of a single organism from video recordings at a large-scale provides means for the quantitative study of its behavior, both individual and collective. This task is particularly difficult for organisms that interact with one another, overlap, and occlude parts of their bodies in the recording. Here we propose WormSwin - an approach to extract single animal postures of Caenorhabditis elegans (*C. elegans*) from recordings of many organisms in a single microscope well. Based on transformer neural network architecture our method segments individual worms across a range of videos and images generated in different labs. Our solutions offers accuracy of 0.990 average precision (AP_0.50_) and comparable results on the benchmark image dataset BBBC010. Finally, it allows to segment challenging overlapping postures of mating worms with an accuracy sufficient to track the organisms with a simple tracking heuristic. An accurate and efficient method for *C. elegans* segmentation opens up new opportunities for studying of its behaviors previously inaccessible due to the difficulty in the worm extraction from the video frames.

## Introduction

Behaviour is the external output of an animal’s nervous system. The possibility to systematically observe, extract, and quantify an animal’s motion is a prerequisite to investigate and ultimately understand its behavioral repertoire. Alterations to an organism’s natural behavior is a phenotypic readout of the neural and other molecular changes that are causing them. To fully understand the functioning of neural mechanisms it is therefore essential to dissect their effect on an animal’s behavior.

Capturing behavior requires video acquisition systems allowing to either view or infer an entire posture of an organism and its change in time. One of the main challenges in obtaining complete and precise posture measurements are the occlusions of animal body parts in a 2D video recording, especially if more than one individual is being imaged. To resolve this, extensive 3D motion capture systems have been developed [1] as well as methods that allow to impute the occluded parts of the posture [2].

These challenges have not yet been resolved for the model organism *C. elegans.* While imaging the nematode’s behavior is less complex than imaging of larger organisms and massively parallel recording systems allow to capture thousands of worms at a time [3,4], there are currently no end-to-end methods that resolve their postures when occlusions occur. The quantification of *C. elegans* strains’ behavior and characterization of their phenotypes is therefore based on segments of worm motion in which it does not coil or intersect with another worm. As a result, a large portion of the worm behavior, including its group behavior, cannot be quantitatively analyzed.

Here we propose an automated method for *C. elegans* posture extraction from 2D video recordings. Based on deep learning transformer architecture and a classical instance segmentation training objective, our solution allows to correctly infer an outline of an individual worm body in overlapping and occluded configurations. We train the neural network on randomly generated image data, obtaining a solution that generalizes to various real datasets. With the segmentation outputs of our method we are able to correctly infer worm trajectories with a simple position matching heuristics. WormSwin opens up new opportunities to study the full repertoire of *C. elegans* behavior including behaviors such as mating that were previously inaccessible to quantitative analysis.

## Related Work

Over the past years different methods for *C. elegans* detection and segmentation have been proposed, either as part of a general approach to tracking and behavioral studies, or as a stand-alone method.

One of the first methods for automated worm tracking and behavior quantification was proposed by Baek et al. [5]. The method used a computer-controlled tracker for single worms, recording grayscale videos. The grayscale frames of a video were binarized based on the mean and standard deviation of pixel intensities and a predefined threshold. The method computes features such as the area of foreground or the movement between two frames in the binarized videos and uses them as input to the algorithm [6] for classification of different *C. elegans* strains. Swierczek et al. [7] proposed a tracking approach called Multi Worm Tracker. The method calculates a background estimate using pixel intensity values. Moving objects are found by searching for pixels darker than the background by a specific threshold. In the next frame, the objects are searched for in the vicinity of their previous location.

The arrival of deep learning offered new opportunities to build more accurate methods for worm segmentation and tracking. Javer et al. [8] developed a multi-object tracking framework able to track *C. elegans* as well as fish and drosophila larvae. The method requires manual tuning of segmentation parameters to best perform with the given recorded data and comes with a graphical user interface for the ease of use and evaluation of the results. Using the motion data, the framework extracts a large number of features characterizing worm movement. Hebert et al. [9] proposed a pose estimation method for videos of single moving *C. elegans* in challenging poses like coiling. Using a ResNetV2 [10]-like architecture the centerline of worms is predicted. With the help of temporal information the head and tail position is determined. Wählby et al. [11] proposed a phenotype analysis toolbox based on the open-source CellProfiler [12] project. To untangle clusters of worms the authors describe them as a mathematical graph and, using a learned model of worm postures, search for the best representation of true worms. The worm posture model is based on a training dataset of isolated single *C. elegans* shapes and on computed angle-based shape descriptors. One of the downsides of this approach is that unexpected phenotypes are likely to be discarded as debris. Banerjee et al. [13] introduced a deep learning *C. elegans* tracking method in which the detection is based on YOLOv5 [14] and tracking on Strong SORT algorithm [15]. For each detected object the method outputs its bounding box, then threshold-based segmentation and skeletonization are applied to infer shapes of the detected objects. Fudickar et al. [16] developed a two-shot segmentation method based on Mask R-CNN [17] with ResNet-101 [18] backbone, to segment *C. elegans* imaged in petri-dishes with a low-cost image capturing system. However, the method did not solve the problem of segmenting overlapping worms and segments them as one object. Mais et al. [19] developed a proposal-free instance segmentation method, called PatchPerPix, based on a convolutional neural network (CNN) trained to predict the shape of a *C. elegans* in a small patch of the whole image (local shape patches). The method uses a modified U-Net [20] deep neural network and patch affinity graphs to reconstruct individual worm shapes. For each pixel the method predicts which shape patch it belongs to and, using a patch affinity graph, merges the patches to form complete instance shapes. Lalit et al. [21] proposed an embedding-based instance segmentation method for 2D and 3D microscopy data, called EmbedSeg. The method is based on ERF-Net [22], predicting spatial embeddings of pixels. These embeddings are then clustered into object instances. To train this method, an additional step of pre-processing the dataset is required to generate object-centered image patches for every object. The method was tested on different datasets including the *C. elegans* BBBC010 dataset.

Among the methods described above there are one- and two-shot detectors. One shot-detector architectures like YOLO [23] detect objects in one step. Pre-defined boxes (also called anchors) are placed onto a grid, laid over the image. For each box, the network predicts if the box contains an object. On the other hand, two-shot detectors consist of a region proposal network (RPN) proposing regions of interest (RoI) to a second network, refining these proposals to form the actual predictions. One-shot object detection methods (like [13]) are in general less computationally expensive compared to two-shot approaches (e.g. [16]), although the latter ones achieve a higher precision especially in more challenging scenes. This is one of the reasons for the high popularity of two-shot networks such as Mask R-CNN in the domain of instance segmentation.

Usually more than one box is predicted per object. To only keep the best matching box, many methods apply non-maximum suppression (NMS). This approach consists of removing from the predicted highly overlapping bounding boxes those with lower probability values as potential false positive detections of the same object. However, NMS can lead to removal of correct detections, especially in dense scenes, where many objects in the image overlap.

In this paper we address the problem of segmenting objects in dense scenes by combining the well established architecture of two-shot detectors with state of the art vision transformer. To avoid the pitfalls of the NMS algorithm, we apply Soft Non Maximum Suppression (Soft-NMS) [24].

## Methods

### Datasets

#### CSB-1 Dataset

The CSB-1 dataset consists of 56 videos with a length of ~ 1.5 min., a frame rate of 5 Hz and frame size of 912 x 736 px which were generated to describe the new *C. elegans* csb-1 strain [25]. We annotated 10 of those videos, where nine videos were reserved for training and one for testing. The videos do not contain any visible petri-dish edges, have different backgrounds and varying numbers of worms. We extracted frames from the videos using *FFmpeg*^1^ and used them to generate our synthetic training dataset described below.

Worms were annotated individually with a binary mask labelling foreground pixels, resulting in one mask image per worm per frame. These separate masks allow to mark all the worms also in cases where pixels of individual *C. elegans* overlap. The labeled CSB-1 dataset contains more than 60,500 individual worm masks. *C. elegans* at the image borders are ignored during the labelling process. Our data is available under ^2^ as a rich resource to develop better methods for animal tracking.

#### Synthetic Dataset

For training the model we generated a synthetic dataset using the nine annotated videos from the CSB-1 dataset described above. We automatically cut out foreground objects from the original grayscale images, according to their polygon annotations and created patches with a worm in the foreground and transparent background. Additionally, we created background images as templates by removing all foreground objects using standard graphics software and filled them with patches of background, taken from the same background images.

The following pipeline was applied to create each image of the synthetic dataset:

1. Randomly select 5 to 30 foreground objects and a background template
2. Randomly flip and rotate foreground objects and their corresponding annotations
3. Apply blurring to foreground objects by averaging the pixel values using a 2×2 px kernel
4. Place foreground object patches on background image:

a. In 20% cases: place a foreground object on top of another one
b. In 80% cases: place a foreground object randomly on the background image

The generated training dataset consists of 10,000 grayscale images with a size of 912 x 736 px and more than 175,000 labeled *C. elegans.* We generated additional 1,000 images for testing. We added grayscale rings of random sizes surrounding the center of the images (see Fig. 1a) to make the network robust against similar artifacts (e.g. petri-dish edges) in other test datasets. Foreground objects might overlap with the artificial petri-dish edges, but are only placed on the inside of the rings. Using the object masks of the original data, for each foreground object we generated a binary mask corresponding to its artificially generated location and shape. These masks are used as ground truth for model training and testing on this dataset. Our synthetic training dataset is available at^3^.

**Fig 1.**
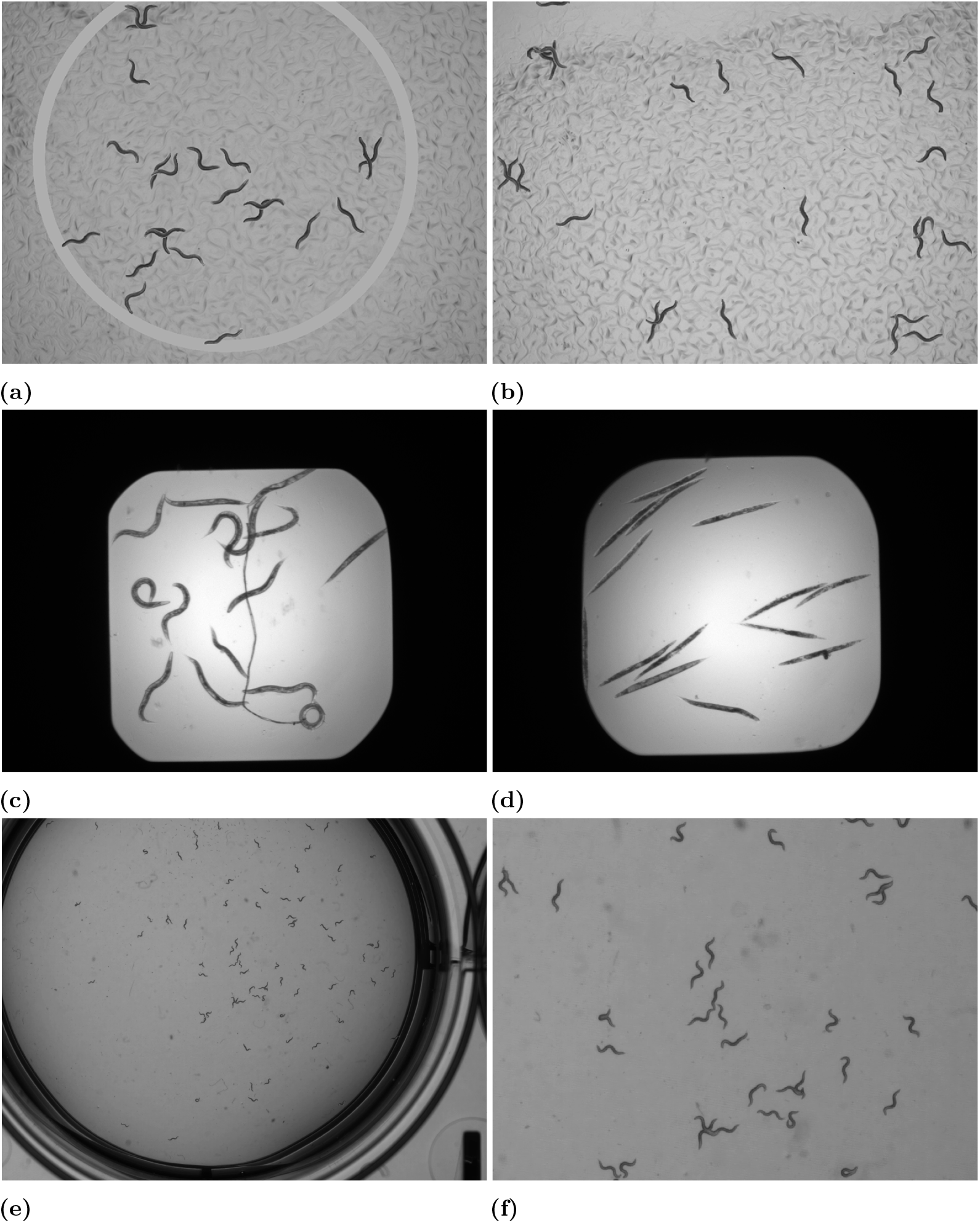
Example images from the datasets used in this study: **(a)** Synthetic Dataset example with added ring, **(b)** Synthetic Dataset without ring, **(c)** BBBC010 Dataset example with mostly alive *C. elegans,* **(d)** BBBC010 Dataset patch with mostly dead *C. elegans,* **(e)** Mating Dataset with petri-dish ring, **(f)** Zoomed-in Mating Dataset patch with many overlaps.

#### BBBC010

The “BBBC010 - *C. elegans* live/dead assay” [26] (BBBC010) dataset consists of 200 images, divided into 100 bright-field and 100 green fluorescent protein (GFP) microscopy images of the same scene. The images have a size of 696 x 520 px and are saved as 16-bit grayscale TIFF files. For our experiments we converted the images to 8-bit grayscale PNG images. The images contain a black border surrounding the region of interest (ROI) with the *C. elegans* in the center (Fig. 1c) which makes up around 50% of the image. Ground truth consists of binary foreground/background images for each worm separately, allowing to disentangle the overlapping shapes.

The images show *C. elegans* exposed to *Enterococcus faecalis* with a negative control group containing dead worms and a positive control group, which was treated with ampicillin and includes alive worms. While the alive *C. elegans* have the natural curved shapes (Fig. 1c), the negative control group appear rod-like with an uneven texture (Fig. 1d).

#### Mating Dataset

The Mating Dataset (MD) was created from a 10 min. long video with a frame rate of 25 Hz and a frame size of 3,036 x 3,036 px. It contains freely moving worms as well as mating ones. Mating behavior is particularly difficult to segment as the two individuals are strongly overlapping and parallel to one another (Fig. 1e). This dataset represents therefore the most challenging segmentation task for our method.

We downsampled the video to 5 Hz and selected 50 frames randomly for annotation and testing of our approach. More than 3,900 individual worm postures were labeled in this dataset. The labeling includes only mature *C. elegans,* worms touching the image boundary were ignored. We split the frames into 450 images with a size of 1,012 x 1,012 px without overlap. The grayscale images show *C. elegans* in a petri-dish with the edges visible (see example patch in Fig. 1f).

## Network architecture

To predict bounding boxes and instance segmentation masks we use the Hybrid Task Cascade (HTC) [27] neural network architecture, combined with Swin Transformer [28] as backbone.

Swin Transformer is a Vision Transformer (ViT)-based backbone architecture [29], which can be applied to different vision-related tasks (e.g. classification, detection or segmentation). Previous ViTs divided the input image into relatively large patches and computed self attention among them. ViTs showed lower computational complexity, but did not account for small details in large images. To tackle this problem Swin Transformer introduced a **S**hifted **Win**dow approach to reduce the computational complexity of standard multi-head self attention (MSA) modules. Additionally, Swin Transformer builds hierarchical feature maps, merging image patches in deeper layers, enabling small-sized patches, leading to more detailed predictions. We chose Swin-L architecture variant in our study which was pre-trained on ImageNet-22K [30] with an image size of 384 x 384 px (similar to [28]).

HTC improves the architecture of Cascade Mask R-CNN [31] by introducing interleaved bounding box regression and instance segmentation mask prediction. The information flow is optimized by adding direct connections between the individual mask branches (Fig. 2). Additionally, a semantic segmentation branch is added to the original architecture to help to distinguish between foreground and background. In our experiments we do not use this additional semantic segmentation branch.

**Fig 2.**
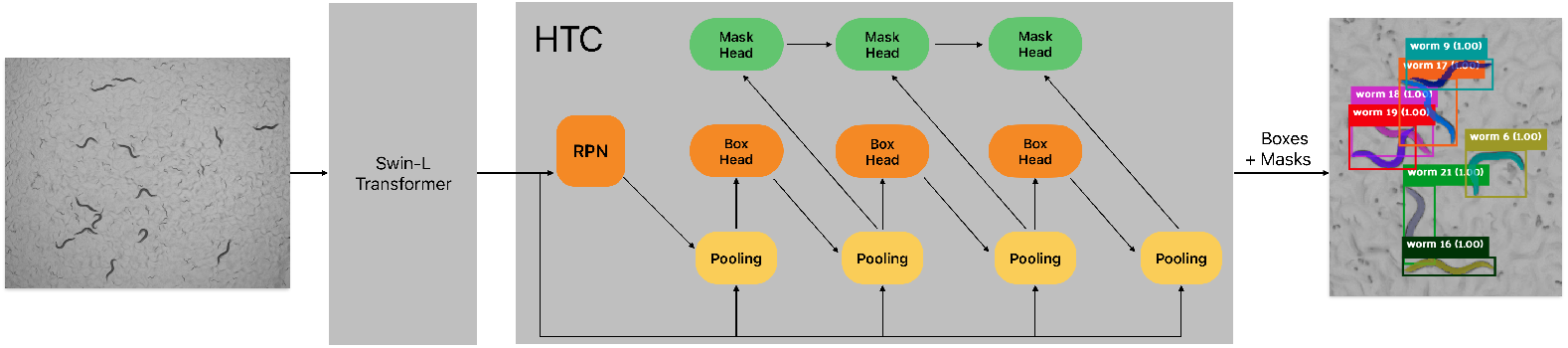
Network Architecture based on Swin-L backbone and HTC. Batch Norm (BN) layers in HTC are replaced by Group Norm (GN) + Weight Standardization (WS). Bounding Box heads are changed from the original Shared2FC architecture to Shared4Conv1FC.

To further improve the accuracy when training on small batches, we exchanged the default Batch Normalization (BN) [32] with Group Normalization (GN) [33] and Weight Standardization (WS) [34] in the HTC. We also replaced the Shared 2 Fully-Connected Bounding Box heads (Shared2FC) by Shared 4 Convolution + one Fully-Connected Bounding Box head (Shared4Conv1FC) [33]. To suppress low quality detections but keep high quality predictions in dense and overlapping scenes we use Soft-NMS instead of the traditional NMS algorithm for the R-CNN during test time (see HTC++ [28]).

### Training

We used multi-scale training with a size between 480 px and 800 px for the shorter side and 1333 px at most for the longer side, AdamW [35] as optimizer, Cosine Annealing Learning Rate Scheduler [36] and Linear Warm-Up [37] (similar to [28]). The learning rate was set to 2.5e^-5^ and weight decay to 0.1. The number of warm-up iterations of the linear warm-up and learning rate scheduler was set to 1,000, warm-up ratio to 0.1 and minimum learning rate ratio to 1e^-5^. During training and testing the NMS threshold for the RPN was set to 0.7, the Soft-NMS of the R-CNN was set to 0.5 during test time.

We used random flipping with a probability of 0.5 and AutoAugment [38] for multiscale resizing and cropping. Additionally, we used the pre-trained weights for the Swin backbone, trained on ImageNet-21K with an image size of 384 x 384 px. We tested our approach on three different datasets: the publicly available BBBC010 dataset, MD and CSB-1 datasets. During testing all images were resized to 800 px on the short side and to no more than 1,333 px on the longer side, preserving the original ratio. We excluded all instances touching image borders as incomplete *C. elegans* instances.

In all our experiments we used the MMDetection framework [39]. Our code and network configuration file for the MMDetection framework are available at^4^.

WormSwin was trained using 4 Nvidia Tesla V100-SMX2 32 GB GPUs, 6 cores of an Intel Xeon Gold 6248 CPU @ 2.50GHz and 100 GB of RAM. With a batch size of four (one image per GPU) and two workers per GPU, training for 36 epochs took ~ 19h. Evaluation on the test set runs at a speed of 2.7 images/s.

## Results

We trained WormSwin on data synthetically generated based on the CSB-1 dataset. The procedure of data generation allows us to control the degree of overlap among individuals and to train the network on a large number of images containing overlapping worms to improve segmentation of dense scenes. Once trained, we evaluated the model on a synthetic test set (see Table 1) as well as on three independent datasets: BBBC010, MD and CSB-1. These datasets come from different labs, show visual variability, and contain different number and degree of overlapping *C. elegans*. We report our results mostly as COCO Average Precision (AP) [40] calculated using pycocotools^5^. For the BBBC010 dataset, we report our results as DSB AP for comparison to other methods. AP is the area under the precision-recall curve and its values are between 0 and 1, with a higher AP representing better performance. Precision and recall of the detection is calculated for instances that show intersection over union (IoU) with the ground truth above a predefined threshold. DSB mAP calculates a mean Jaccard Index by using different IoU thresholds. COCO mAP uses a more complex approach: detections are sorted by descending confidence score. The calculation iterates over all detections in this order, marks them as True Positive (TP) or False Positive (FP) and adds them to calculate the precision until a recall of 1.0 is reached or iterated over all detections. Different IoU thresholds are used to label detections as TP or FP.

**Table 1.** Test results on all instances. “box” and “mask” refer to the accuracy of detection of the bounding box and segmentation mask, respectively. PatchPerPix ppp+dec refers to the network variant, introduced by [19]. (*multi-scale testing, †additional training data)

|  | AP<sub>0.50</sub> | AP<sub>0.75</sub> | mAP<sub>0.50:0.95</sub> |
| --- | --- | --- | --- |
| **CSB-1** |  |  |  |
| WormSwin (box) | 0.990 | 0.976 | 0.819 |
| WormSwin (mask) | 0.990 | 0.675 | 0.585 |
| **Synthetic** |  |  |  |
| WormSwin (box) | 0.989 | 0.978 | 0.909 |
| WormSwin (mask) | 0.977 | 0.918 | 0.679 |
| **BBBC010** |  |  |  |
| PatchPerPix ppp+dec (mask) | 0.939 | 0.891 | 0.775 |
| WormSwin (box)† | 0.985 | 0.949 | 0.823 |
| WormSwin (mask)† | 0.954 | 0.801 | 0.622 |
| WormSwin (mask)*, † | 0.964 | 0.815 | 0.629 |
| **MD** |  |  |  |
| WormSwin (box)† | 0.990 | 0.968 | 0.832 |
| WormSwin (mask)† | 0.980 | 0.551 | 0.542 |

We report our results mostly for two IoU thresholds: 0.5 and 0.75 as well as a mean AP (mAP) for thresholds from 0.5 to 0.95 with a step size of 0.05. One of the most challenging parts for instance segmentation of *C. elegans*, as well as other biological systems, are overlapping objects in dense configurations. To measure the accuracy of our approach explicitly for overlapping objects, we added a dedicated AP metric. We defined overlapping objects as those whose ground truth bounding boxes overlap by more than 25% or whose segmentation masks that any overlap (>0% IoU). We report the AP for all objects as well as for the overlapping objects separately (Table 2).

**Table 2.** Test results for overlapping worms only (* multi-scale testing, †additional training data)

|  | AP<sub>0.50</sub> | AP<sub>0.75</sub> | mAP<sub>0.50:0.95</sub> |
| --- | --- | --- | --- |
| **CSB-1** |  |  |  |
| WormSwin (box) | 0.883 | 0.643 | 0.551 |
| WormSwin (mask) | 0.975 | 0.409 | 0.527 |
| **Synthetic CSB-1** |  |  |  |
| WormSwin (box) | 0.983 | 0.958 | 0.853 |
| WormSwin (mask) | 0.959 | 0.821 | 0.613 |
| **BBBC010** |  |  |  |
| WormSwin (box) † | 0.911 | 0.821 | 0.661 |
| WormSwin (mask) † | 0.873 | 0.565 | 0.488 |
| WormSwin (mask)*, † | 0.895 | 0.573 | 0.501 |
| **MD** |  |  |  |
| WormSwin (box)† | 0.822 | 0.633 | 0.505 |
| WormSwin (mask)† | 0.893 | 0.079 | 0.355 |

### CSB-1 dataset

Although trained on synthetically generated data, our method generalizes fairly well to the real video data with a mAP of 0.819 and 0.585 for the bounding box and mask respectively, lower by only ~ 0.09 mAP compared to the synthetic data. The same metric on the overlapping worms in the CSB-1 dataset are 0.551 and 0.527. While the mAP is lower for the overlapping *C. elegans* compared to the results on the entire dataset, the AP_0.50_ of the bounding box and mask on the overlapping worms are 0.883 and 0.975, respectively. This result suggests that the worms are detected correctly in principle but there exist errors in mask prediction. What these mask prediction errors are, is however not clear at a first glance. Despite the difference between the AP_0.50_ and AP_0.75_ in the overlapping worms, we found that the segmentation masks align in general well with the ground truth (Fig. 3), however pixels on the edges of each object tend to be imprecisely segmented. Due to the small size of a worm mask with ~ 500 px, errors at the edges of the predicted masks represent ~ 30% of all foreground pixels.

**Fig 3.**
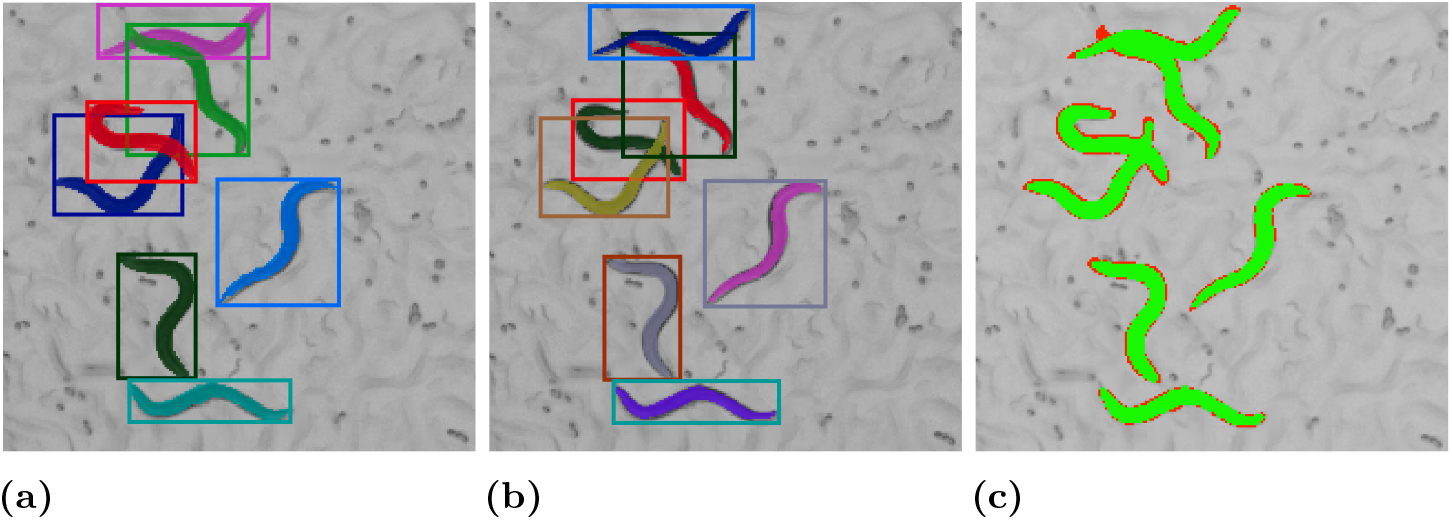
Example from the CSB-1 dataset (box and mask colors are selected randomly). **(a)** Ground truth annotations, **(b)** Predicted bounding boxes and masks, **(c)** TP (green), FP and FN (red) pixels.

To test the hypothesis that most error occur on the mask edges, we implemented an alternative version of the IoU: if a pixel in either ground-truth or predicted mask is at the border of an object (when the 4-way neighborhood is not fully foreground) then it is set to the value of the pixel at this position in the other mask. This way, object border pixels which otherwise would be considered as false negative (FN) or false positive (FP) do not influence the IoU calculation in a negative way. Using this calculation, the mean IoU on the test subset raised from 0.827 to 0.961 (+13.4% increase) on the CSB-1 dataset.

### BBBC010 dataset

Because of the very limited number of training samples (50 images) the predictions of the network trained on BBBC010 were of poor quality. Therefore, we used the network pre-trained on our synthetic data and fine-tune it on 50 randomly selected images from the BBBC010 dataset. We compared the performance of our approach to two existing methods: PatchPerPix [19] and EmbedSeg [21]. To enable this comparison, instead of the COCO AP metric (see Table 1) we used (Data Science Bowl) DSB AP ^6^ as accuracy evaluation on this dataset which was used in the original EmbedSeg method publication [21] (see Table 3).

**Table 3.** Test results using DSB metric (* multi-scale testing, †additional training data, ‡alternative IoU without object edges, mAP for IoUs in range 0.5 to 0.95, step size 0.05)

|  | AP<sub>0.50</sub> | AP<sub>0.60</sub> | AP<sub>0.70</sub> | AP<sub>0.80</sub> | AP<sub>0.90</sub> | mAP |
| --- | --- | --- | --- | --- | --- | --- |
| **BBBC010** |  |  |  |  |  |  |
| PatchPerPix ppp+dec [19] | 0.930 | 0.905 | 0.879 | 0.792 | 0.386 | 0.727 |
| EmbedSeg [21] | 0.965 | 0.934 | 0.896 | 0.762 | 0.326 | - |
| WormSwin (mask)*, † | 0.917 | 0.884 | 0.785 | 0.336 | 0.005 | 0.536 |
| WormSwin (mask)*, †, ‡ | 0.929 | 0.920 | 0.890 | 0.823 | 0.483 | 0.769 |

We used the alternative IoU calculation already used for the CSB-1 dataset, to calculate the DSB accuracy without considering object edges. With the IoU defined this way, using the DSB metric we achieve 0.769 mAP (+0.233), 0.929 AP_0.50_ (+0.012) and 0.823 AP_0.80_ (+0.487) (compare to Tab. 3).

### Mating Dataset

Finally, we tested WormSwin on the MD dataset using weights pre-trained on our synthetic dataset. In this dataset we annotated 50 images, which are larger in size and contain a higher number of *C. elegans* compared to the BBBC010 dataset. Further, we split them into patches of size 1,024 x 1,024 px. We report our results in Table 1). Despite the challenging configurations of worms in this dataset, our method correctly identifies the segmented objects, as indicated by the AP_0.50_ which is comparable to the AP_0.50_ in other datasets. However the AP_0.75_ and mAP_0.50:0.95_ suggest that, while correctly detected, the segmentation masks of the detected objects are imprecise. Similar to other datasets, we hypothesise that these errors occur on the boundaries of the segmentation masks (Fig. 4) as well as are due to the very challenging object overlaps in this dataset.

**Fig 4.**
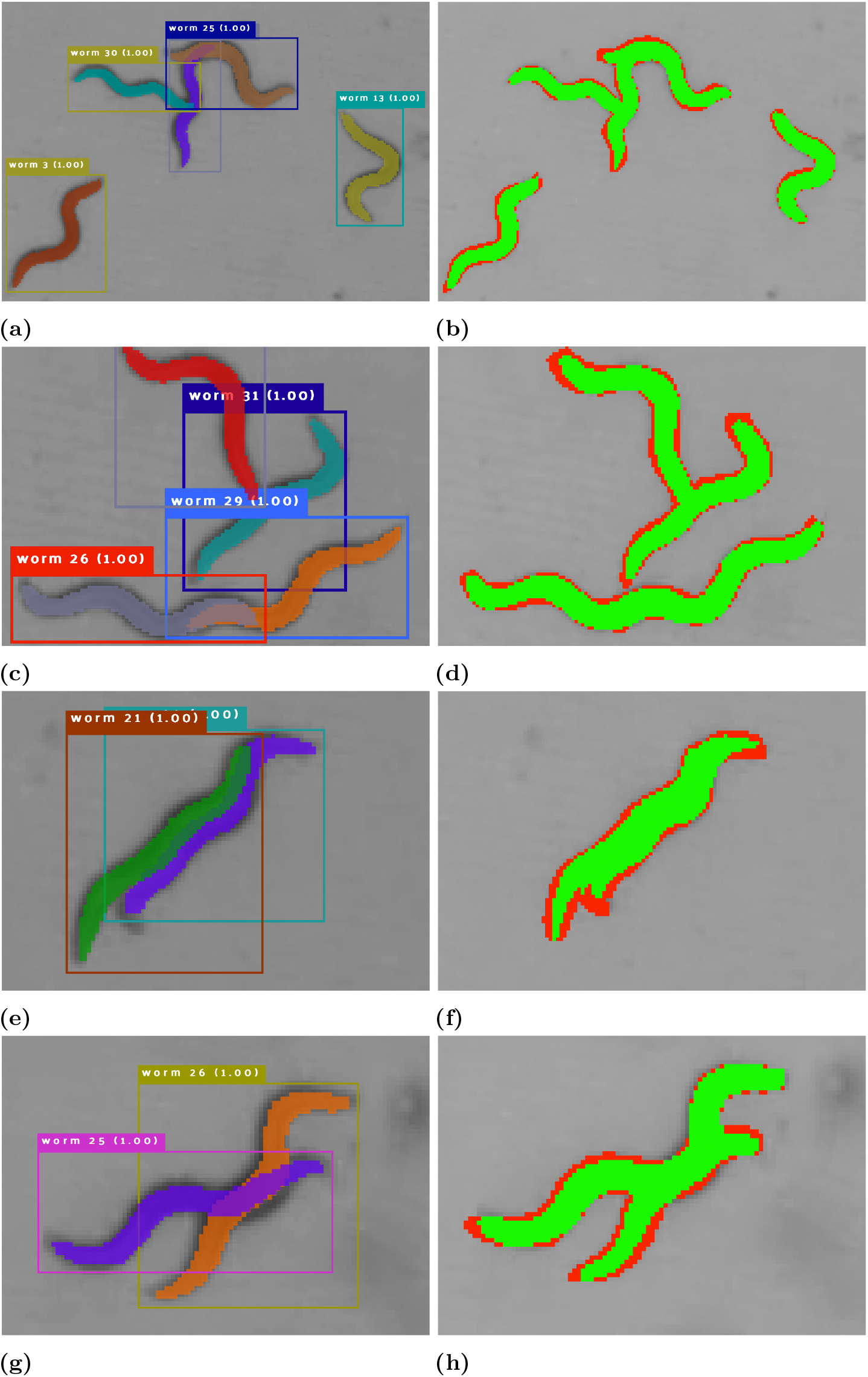
Results on the Mating Dataset (box and mask colors are selected randomly). **(a, c, e, g)** Segmentation results, **(b, d, f, h)** TP (green), FP and FN (red) pixels.

### Tracking

To test if our segmentation results are sufficiently accurate to allow for worm tracking and further behavioral analysis, we implemented a simple IoU-based matching method (Fig. 5) and applied it on our predicted instance segmentation masks in the CBS-1 test set. Between two consecutive frames, objects with the highest overlap in mask are matched into a trajectory. Iterating the matching procedure over all video frames results in object trajectories. In this simple approach, if an object is not detected in a frame but detected in a subsequent frame its trajectory is disrupted and two separate trajectories are created instead. We attempt to reconnect such trajectories in a post-processing step: for 10 frames after loosing an object, starting points of new trajectories are compared with the endpoint of the lost trajectory. If the segmentation masks at these points overlap with at least 50%, the trajectories are reconnected. In the frames with missing segmentation masks the positions of *C. elegans* can be interpolated between two ends of reconnected trajectories.

**Fig 5.**
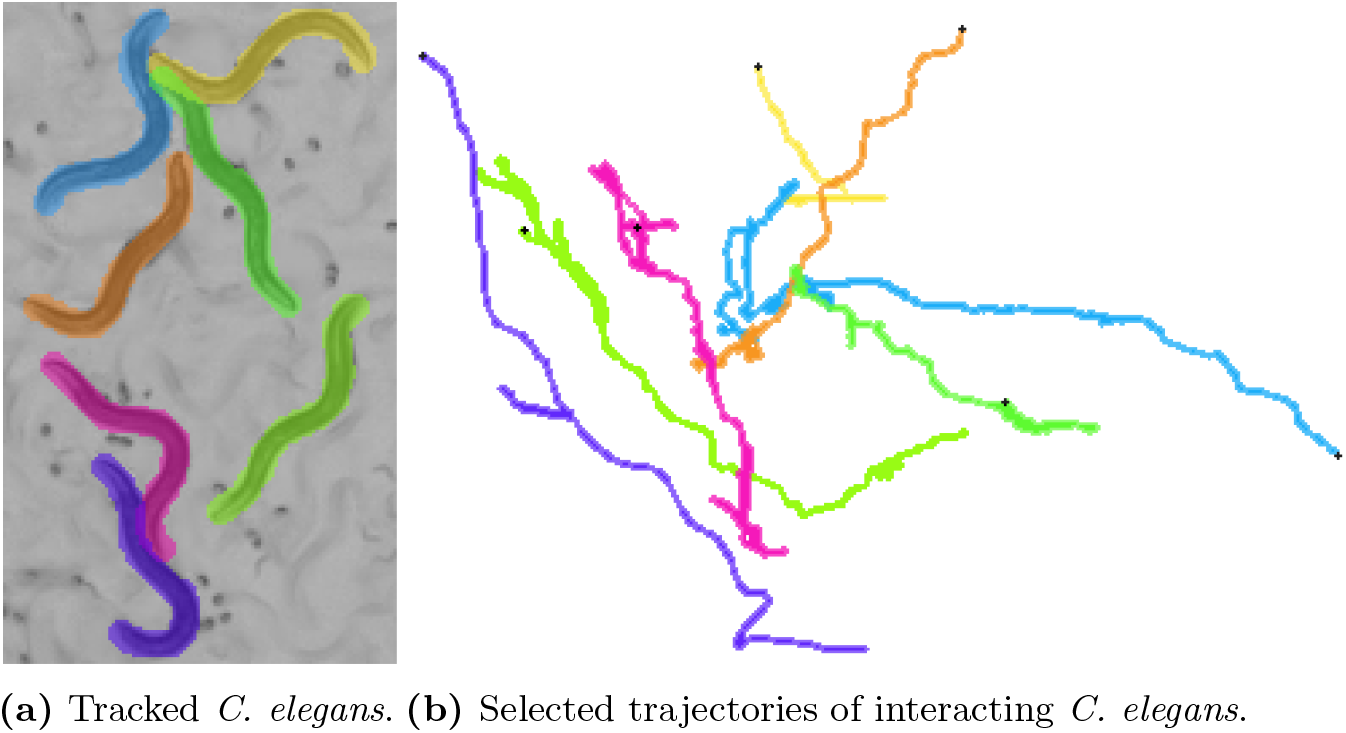
Example of tracked *C. elegans.*

While a tracking method is outside of the scope of this study, our simple approach allows to build trajectories of interacting mating worms (Fig. 5). Tracking these challenging *C. elegans* interactions opens up new possibilities of studying its behavior.

## Discussion

In this work we present WormSwin, a deep learning approach for instance segmentation of microscopy images of *C. elegans*. Our method combines several recent improvements in deep learning and instance segmentation (Transformer Networks, HTC, Group Normalization, Weight Standardization, Soft-NMS) into a single approach trained end-to-end. WormSwin does not require any pre-processing of the image data, enabling researchers to directly apply it on their video or image data.

Together with our method we provide a large dataset of *C. elegans* images with instance mask annotations to help researchers develop better segmentation approaches in the future. The new dataset is by an order of magnitude larger compared to the BBBC010 dataset, enabling training deeper network architectures.

The small size of the BBBC010 benchmark dataset is a limiting factor to extensively train and test our method on this dataset. The accuracy of our method is lower on this dataset compared to the CSB-1 which might be attributed to the differences in the color intensities, size and appearance of *C. elegans* between the two datasets. Since retraining of WormSwin on a small amount of BBBC010 images improved the methods performance, we suggest that to accurately segment datasets differing from CSB-1 characteristics, a similar retraining is necessary.

Notably, our method shows a decrease in AP in the higher IoU threshold categories (e.g. Table 3 AP_0.80_). Despite this precision drop, the segmentation masks appear overall correct (Figures 3–4). We therefore hypothesize that the major errors in the segmentation masks occur on the boundaries of the foreground area and further substantiate this by calculating accuracy metric that does not take into account boundary pixels. The reason for this type of error might be e.g. variation in human-generated labeling. We introduce blurring in the synthetic training data which might additionally change the appearance of the object contours. Despite these errors, individual *C. elegans* poses are captured by the predicted segmentation masks and can be subject to further quantitative analysis.

As a major future improvement of this work we see models exploring temporal information to improve segmentation of overlapping objects. Information on how *C. elegans* individuals arrive in a specific configuration is of great help in disentangling their postures. Such method would additionally combine segmentation with tracking in a single training objective. While generating training datasets for multi-object tracking is a massive work burden, the accuracy of our segmentation approach allows to build preliminary trajectories in an automated fashion.

## Acknowledgments

We would like to thank Matthias Rieckher for supplying us the videos of the CSB-1 dataset, as well as Xiao-Liu Chu for supplying the Mating Dataset video. We thank everyone who helped us annotating the data used in this publication. Maurice Deserno and Katarzyna Bozek were supported by the North Rhine-Westphalia return program (311-8.03.03.02-147635), BMBF program Junior Group Consortia in Systems Medicine (01ZX1917B) and hosted by the Center for Molecular Medicine Cologne. We thank the Regional Computing Center of the University of Cologne (RRZK) for providing computing time on the DFG-funded (Funding number: INST 216/512/1FUGG) High Performance Computing (HPC) system CHEOPS as well as support.

## Footnotes

1 https://ffmpeg.org

2 https://doi.org/10.5281/zenodo.7456803

3 https://doi.org/10.5281/zenodo.7456803

4 https://github.com/bozeklab/worm-swin

5 https://github.com/cocodataset/cocoapi

6 https://www.kaggle.com/competitions/data-science-bowl-2018/overview/evaluation

